# The green autofluorescence intensity and asymmetry of older men is significantly higher than those of older women in their fingernails and certain regions of skin

**DOI:** 10.1101/509562

**Authors:** Yue Tao, Mingchao Zhang, Danhong Wu, Yujia Li, Weihai Ying

**Affiliations:** Med-X Research Institute and School of Biomedical Engineering, Shanghai Jiao Tong University, Shanghai 200030, P.R. China; Department of Neurology, Shanghai Fifth People’s Hospital, Fudan University, Shanghai, P.R. China; Collaborative Innovation Center for Genetics and Development, Shanghai 200043, P.R. China

**Author notes:** These authors contributed equally to this work (Yue Tao, Mingchao Zhang). Corresponding authors: Weihai Ying, Ph.D., Professor, School of Biomedical Engineering and Med-X Research Institute, Shanghai Jiao Tong University Shanghai, 200030, P.R. China.

**Keywords:** Autofluorescence, Gender, Skin, Fingernails, Older people

## Abstract

Our recent studies have suggested that the patients of multiple diseases have characteristic Pattern of Autofluorescence (AF) in their skin and fingernails, which may become novel biomarkers for both disease diagnosis and evaluation of health state. Since male populations may have higher levels of oxidative stress and inflammation than female population, in our current study we tested our hypothesis that the green AF intensity of older men is higher than that of older women in their fingernails and skin. We found that in both left and right Index Fingernails, the green AF intensity of the men of both the age group of 61 - 70 years of old and the age group of 71 - 80 years of old is significantly higher than that of the women of the same age groups. At both left Dorsal Centremetacarpus and left Centremetacarpus, the green AF intensity of the men at the age between 71 - 80 years of old is also significantly higher than that of the women of the same age group. Moreover, in Index Fingernails, Dorsal Centremetacarpus and Centremetacarpus, the green AF asymmetry of the older men of certain age groups is significantly higher than that of the women of the same age groups. Collectively, our study has provided the first evidence indicating the gender difference between the green AF intensity and asymmetry of older men and those of older women in their fingernails and certain regions of skin, which is valuable for establishing the AF-based diagnostic method.

## Introduction

Our recent studies have suggested that characteristic ‘Pattern of Autofluorescence (AF)’ could be a novel biomarker for non-invasive diagnosis of multiple major diseases, including acute ischemic stroke (AIS) (3), myocardial infarction (MI) (17), stable coronary artery disease (17), Parkinson’s disease (4) and lung cancer (13). Our study has also indicated that the AF intensity is highly correlated with the risk of developing into AIS, suggesting that the green AF may be used for evaluating a person’s state of cardiovascular health (12).

Our studies have suggested that oxidative stress is a key factor that can produce the increased epidermal green AF by altering keratin 1 (10,11). Our latest study has indicated that inflammation can also induce increased green AF of mouse’s skin (19). Because there are studies suggesting that compared with older men, older women have longer life expectancy (9,15) and lower levels of oxidative stress (2,6–8,16) and inflammation (1), we proposed our hypothesis that the green AF intensity of older men is significantly higher than that of older men in their fingernails and certain regions of skin. In current study, we determined the green AF intensity and asymmetry of the fingernails and certain regions of the skin of both elderly men and women. Our observations have provided evidence supporting our hypothesis.

## Methods and materials

### Human subjects

The human subjects have lived in three communities of Shanghai, P.R. China. The human subjects in our study were divided into three groups: Group 1: The Age Group between 61 - 70 years of old; Group 2: The Age Group between 71 - 80 years of old; and Group 3: The Age Group between 81 - 90 years of old.

### Determinations of the Autofluorescence of Skin and Fingernails

A portable AF imaging equipment was used to detect the AF of the fingernails and certain regions of the skin of the human subjects. The excitation wavelength is 485 nm, and the emission wavelength is 500 - 550 nm. For all of the human subjects, the AF intensity in the following seven regions on both hands, i.e., fourteen regions in total, was determined, including the Dorsal Index Finger, Ventroforefingers, Dorsal Centremetacarpus, Centremetacarpus, Dorsal Antebrachium, Ventribrachium and Index Fingernails.

### Statistical analyses

All data are presented as mean + SEM. Data were assessed by Student *t* test. *P* values less than 0.05 were considered statistically significant.

## Results

### 1. The green autofluorescence intensity of the men at the age between 61 - 80 years of old is significantly higher than that of the women of the same age group in their fingernails

In both left and right Index Fingernails, the green AF intensity of the men at the age between 61 - 70 years of old is significantly higher than that of the women of the same age group (Fig. 1). The green AF intensity of the men at the age between 71 - 80 years of old is also significantly higher than that of the women of the same age group (Fig. 1). We further found that the green AF intensity of the women at the age between 81 - 90 years of old is significantly higher than that of both the women at the age between 61 - 70 years of old and the women at the age between 71 - 80 years of old in left Index Fingernails (Fig. 1).

**Fig. 1.**
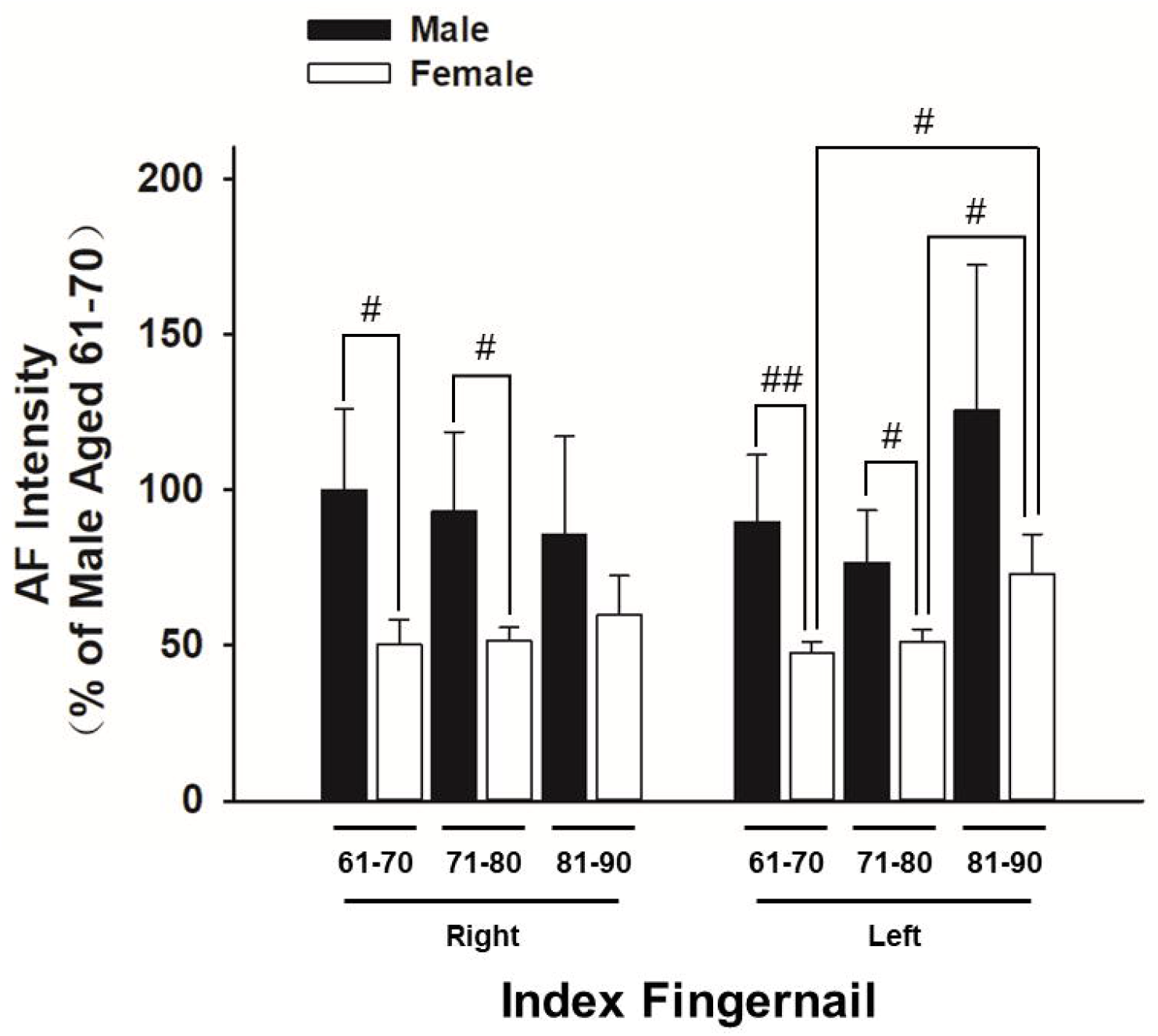
The green autofluorescence intensity of the men at the age between 61 – 80 years of old is significantly higher than that of the women of the same age group in their fingernails. In both left and right Index Fingernails, the green AF intensity of the men at the age between 61 - 70 years of old is significantly higher than that of the women of the same age group. The green AF intensity of the men at the age between 71 - 80 years of old is also significantly higher than that of the women of the same age group. #, *P* < 0.05 (*t*-test); ##, *P* < 0.01 (*t*-test). The number of the men in the age group of 61 - 70, 71 - 80, and 81 - 90 years of old is 25, 20 and 11, respectively. The number of the women in the age group of 61 - 70, 71 - 80, and 81 - 90 years of old is 64, 48 and 15, respectively.

### 2. The green autofluorescence intensity of the men at the age between 71 - 80 years of old is significantly higher than that of the women of the same age group in certain regions of skin

At left Dorsal Centremetacarpus, the green AF intensity of the men at the age between 71 - 80 years of old is significantly higher than that of the women of the same age group (Fig. 2A). At left Centremetacarpus, we also found a similar difference between that green AF intensity of the men at the age between 71 - 80 years of old and that of the women of the same age group (Fig. 2B). At other regions examined, including Dorsal Index Finger (Fig. 2C), Dorsal Antebrachium (Fig. 2D), Ventroforefinger (Fig. 2E) and Antebrachium (Fig. 2F), there was no significant differences between the green AF intensity of the older men and that of the women of the same age group.

**Fig. 2.**
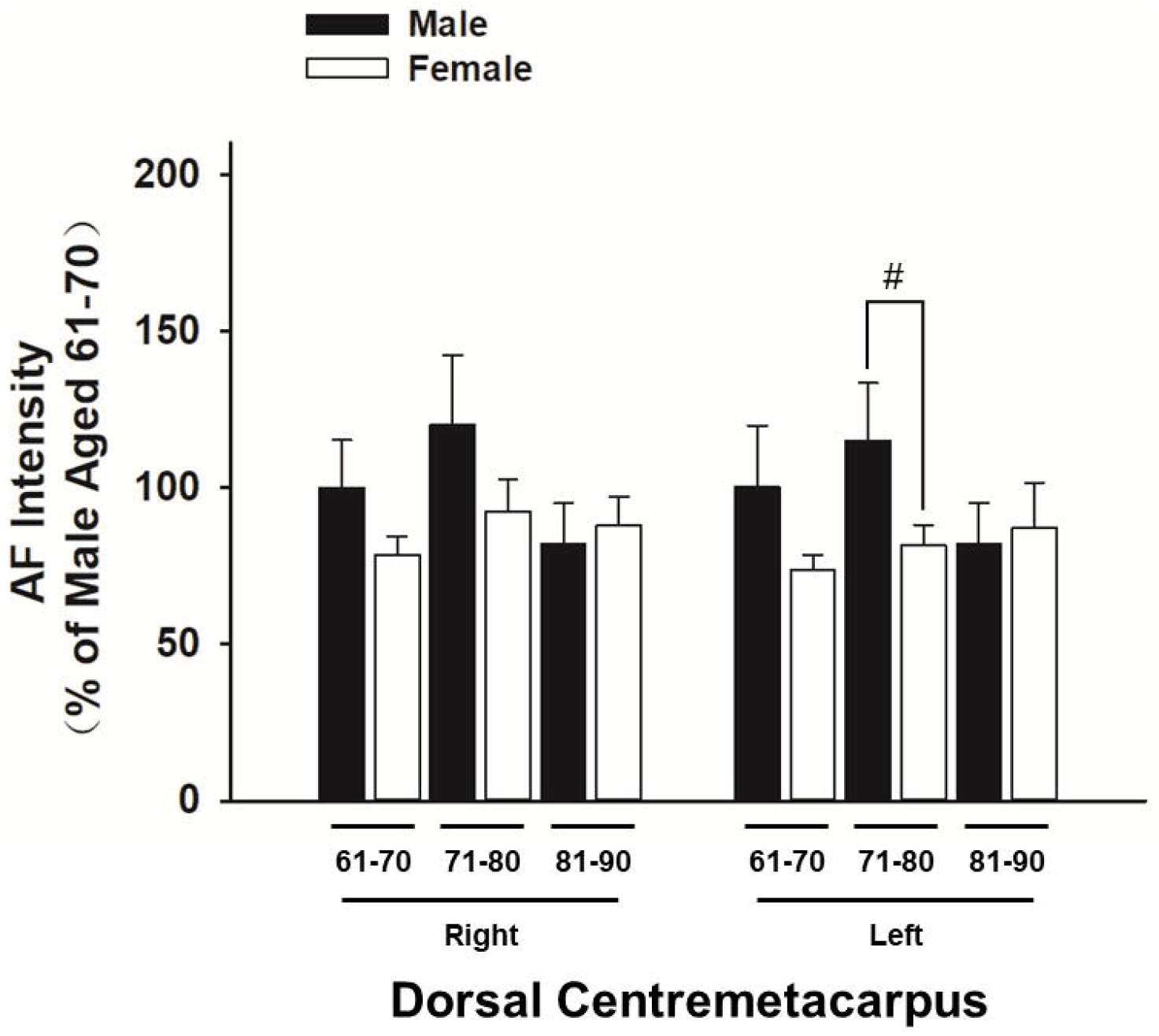
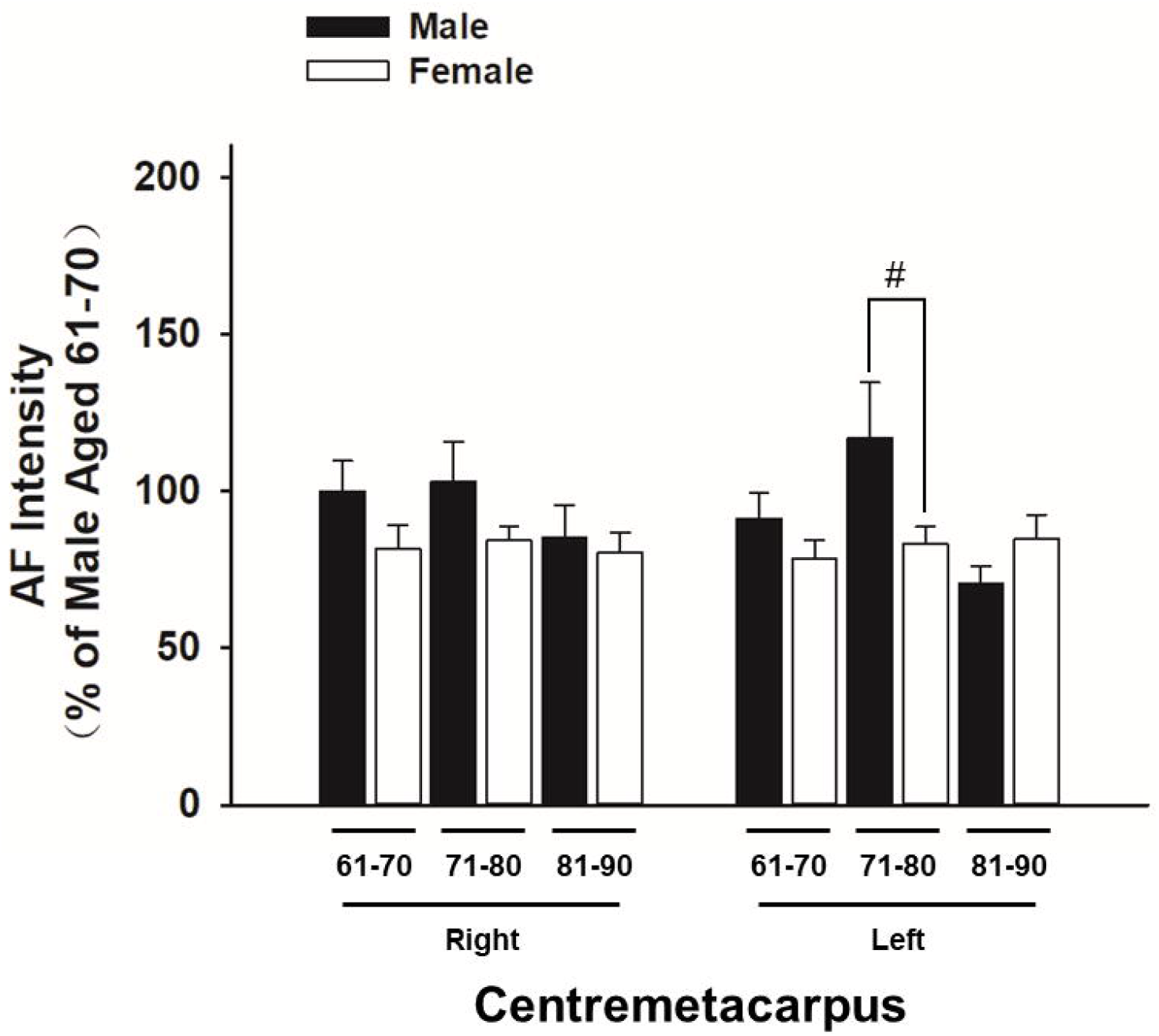
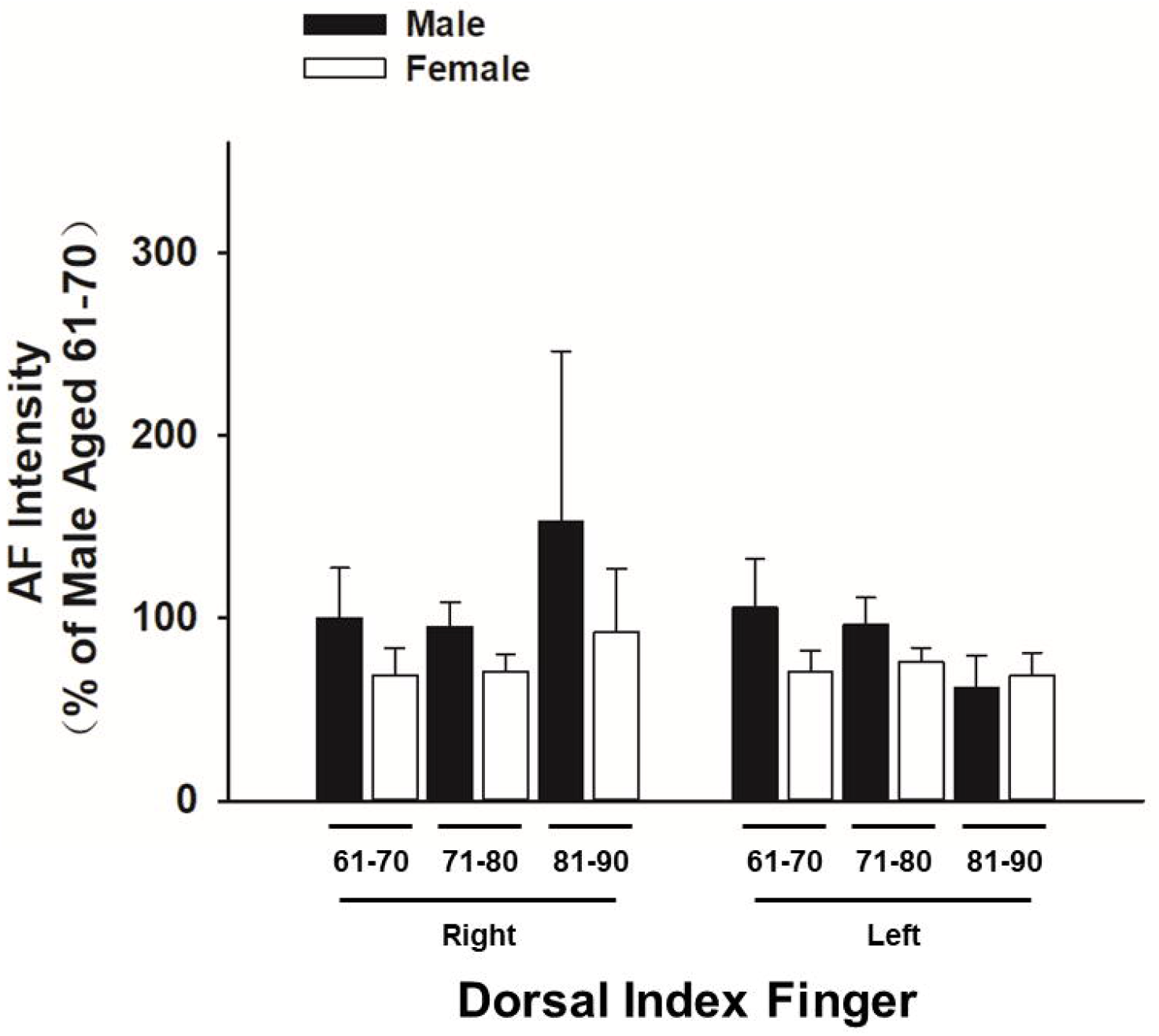
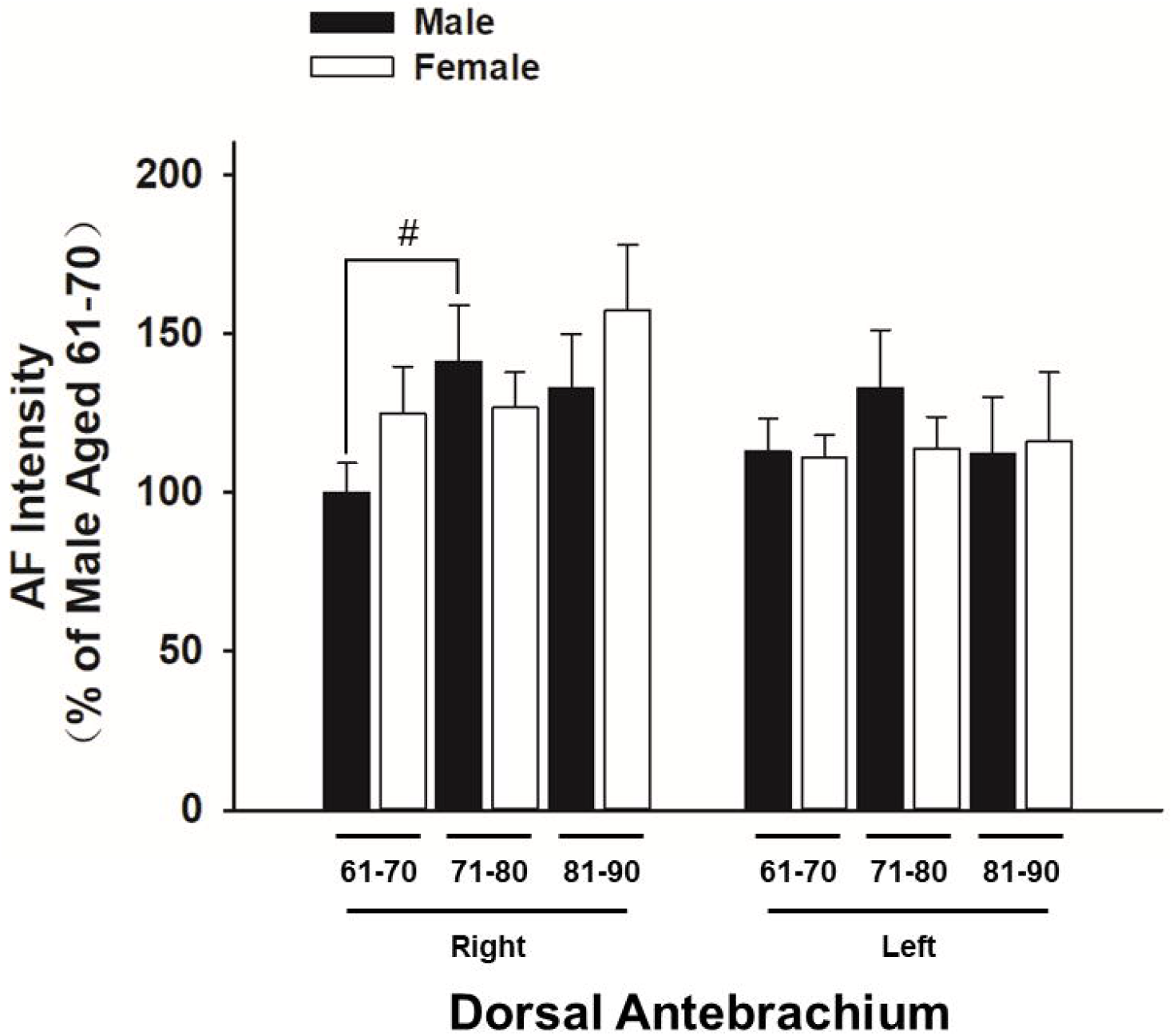
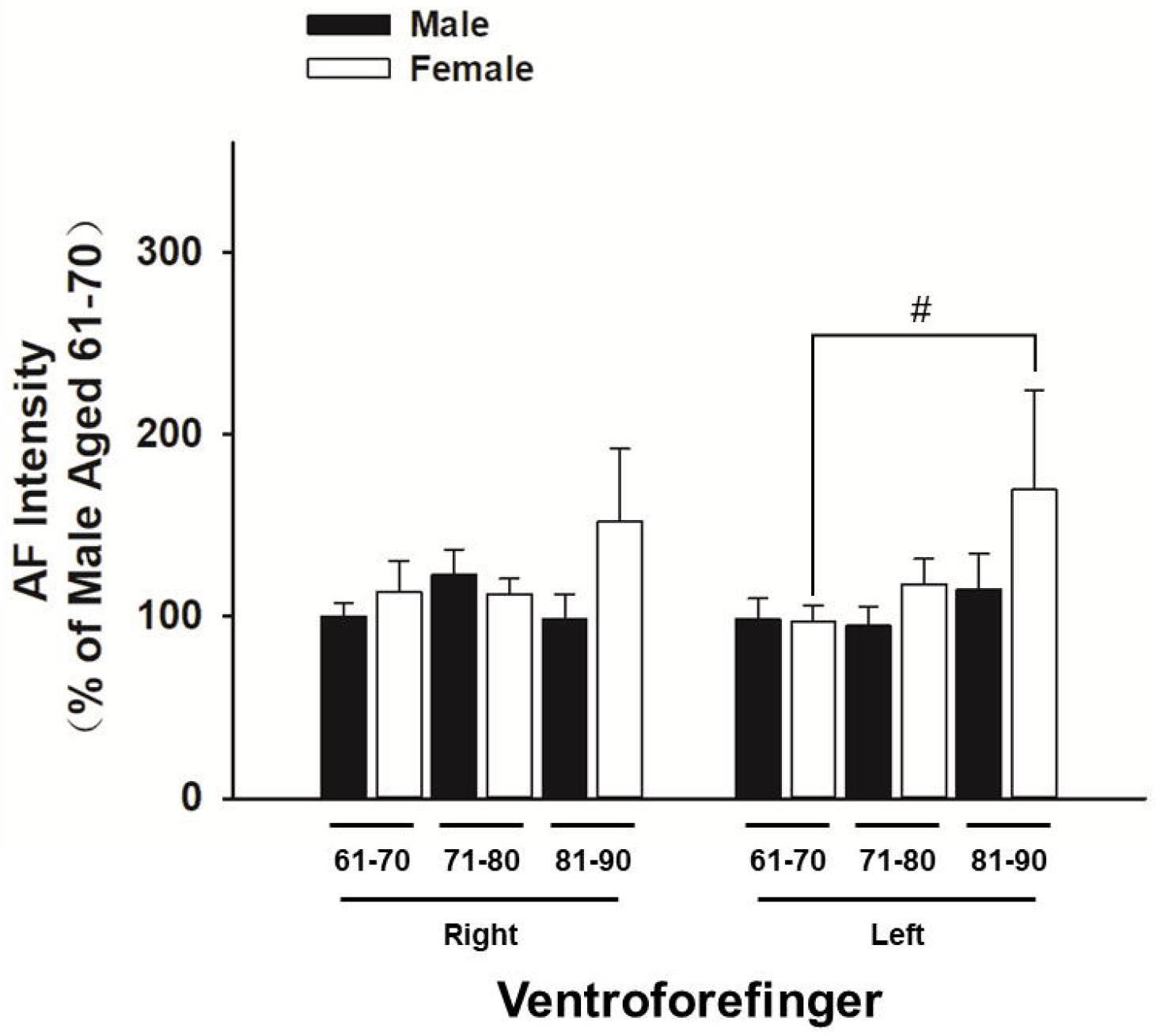
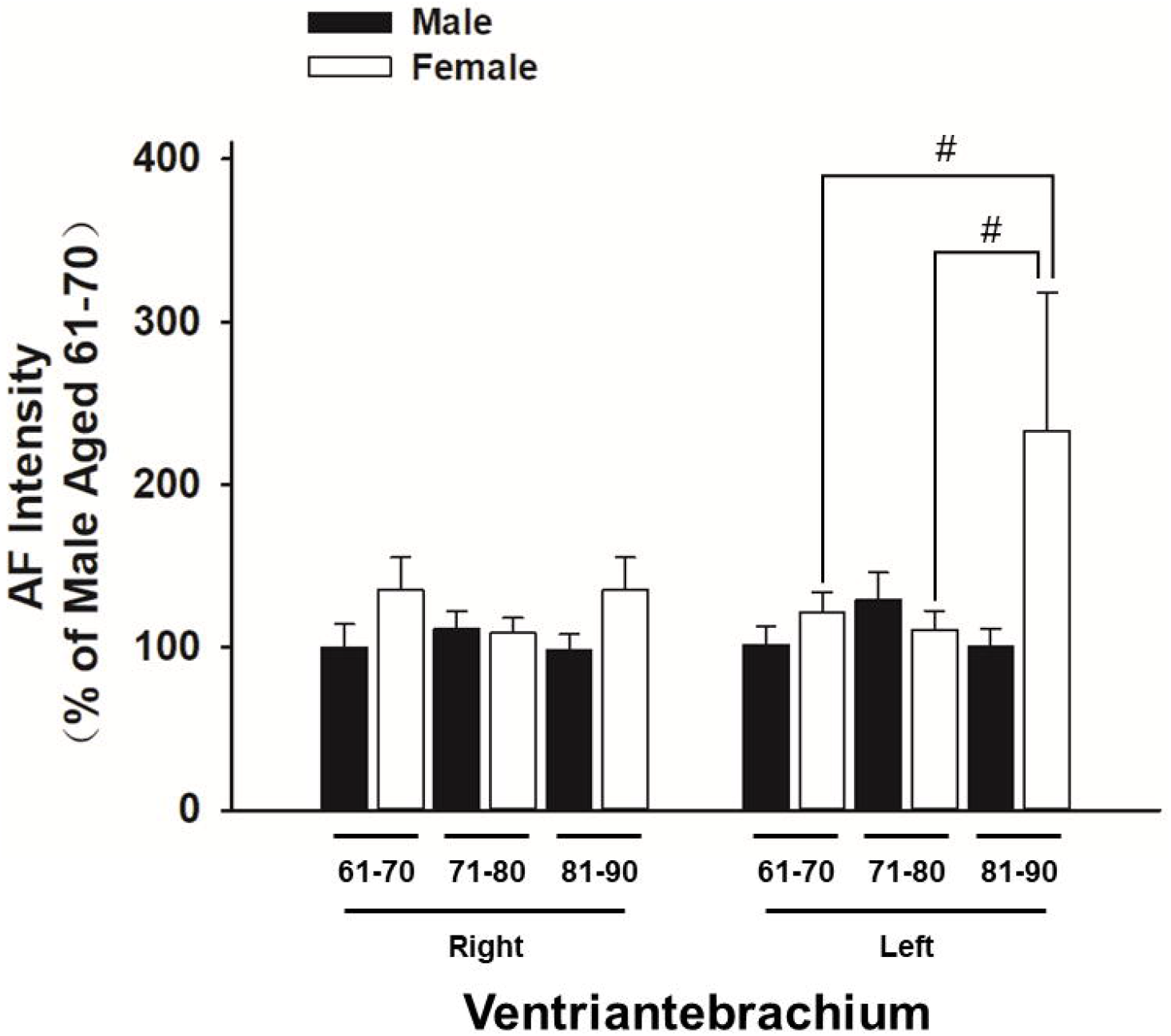
The green autofluorescence intensity of the men at the age between 71 – 80 years of old is significantly higher than that of the women of the same age group in certain regions of skin. (A) At left Dorsal Centremetacarpus, the green AF intensity of the men at the age between 71 – 80 years of old is significantly higher than that of the women of the same age group. (B) At left Centremetacarpus, there was a similar difference between the green AF intensity of the men at the age between 71 - 80 years of old and that of the women of the same age group. At other regions examined, including Dorsal Index Finger (2C), Dorsal Antebrachium (2D), Ventroforefinger (2E) and Antebrachium (2F), there was no significant differences between the green AF intensity of the older men and that of the women of the same age group. #, *P* < 0.05 (*t*-test). The number of the men in the age group of 61 - 70, 71 - 80, and 81 - 90 years of old is 25, 20 and 11, respectively. The number of the women in the age group of 61 - 70, 71 - 80, and 81 - 90 years of old is 64, 48 and 15, respectively.

We further found that the green AF intensity of the women at the age between 81 - 90 years of old is significantly higher than that of the women at the age between 61 - 70 years of old at left Ventroforefinger (Fig. 2E). At left Ventriantebrachium, the green AF intensity of the women at the age between 81 - 90 years of old is significantly higher than that of both the women at the age between 61 - 70 years of old and the women at the age between 71 - 80 years of old (Fig. 2F).

### 3. The green autofluorescence asymmetry of the older men is not significantly different from that of the women of the same age groups in their fingernails

In Index Fingernails, the green AF asymmetry of the men at the age between 61 - 70 years of old shows the tendency to be higher than that of the women of the same age group, which did not reach statistical significance (Fig. 3). There was a similar difference between the green AF asymmetry of the men at the age between 81 - 90 years of old and that of the women of the same age group (Fig. 3).

**Fig. 3.**
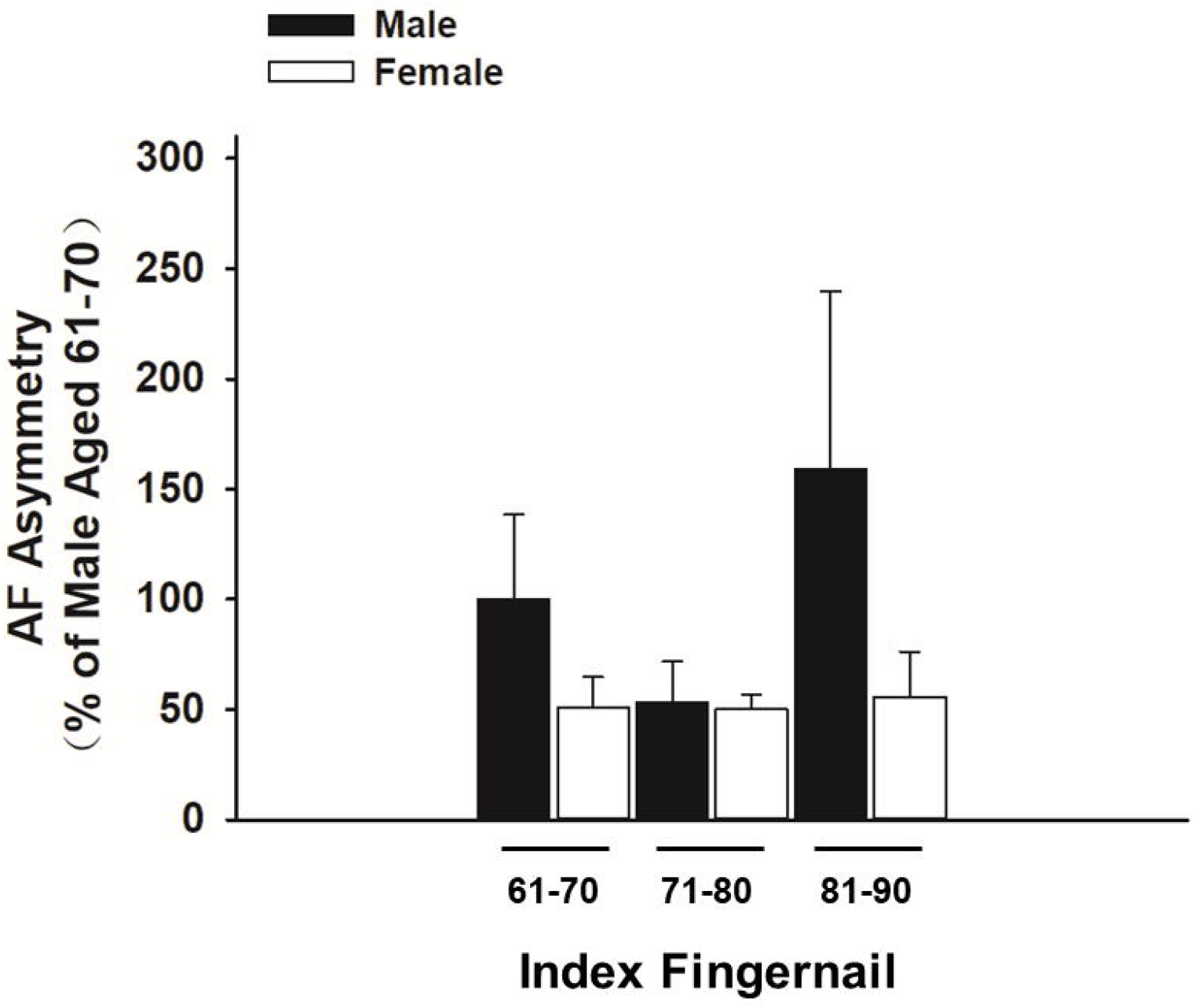
The green autofluorescence asymmetry of the men at the age between 61 – 90 years of old is not significantly different from that of the women of the same age group in their fingernails. In Index Fingernails, the green AF asymmetry of the men at the age between 61 - 70 years of old shows the tendency to be higher than that of the women of the same age group, which did not reach statistical significance. There was a similar difference between the green AF asymmetry of the men at the age between 81 - 90 years of old and that of the women of the same age group. The number of the men in the age group of 61 - 70, 71 - 80, and 81 - 90 years of old is 25, 20 and 11, respectively. The number of the women in the age group of 61 - 70, 71 - 80, and 81 - 90 years of old is 64, 48 and 15, respectively.

### 4. The green autofluorescence asymmetry of the men at the age between 71 - 80 years of old is significantly higher than that of the women of the same age group in certain regions of skin

At Dorsal Centremetacarpus, the green AF asymmetry of the men at the age between 71 - 80 years of old is significantly higher than that of the women of the same age group (Fig. 4A). At left Centremetacarpus, we also found a similar difference between the green AF asymmetry of the men at the age between 71 - 80 years of old and that of the women of the same age group (Fig. 4B). At other regions examined, including Dorsal Index Finger (Fig. 4C), Dorsal Antebrachium (Fig. 4D), Ventroforefinger (Fig. 4E) and Antebrachium (Fig. 4F), there was no significant differences between the green AF intensity of the older men and that of the women of the same age group. We further found that the AF asymmetry of the women at the age between 81 - 90 years of old is significantly higher than that of the women at the age between 71 - 80 years of old at Ventroforefinger (Fig. 4E).

**Fig. 4.**
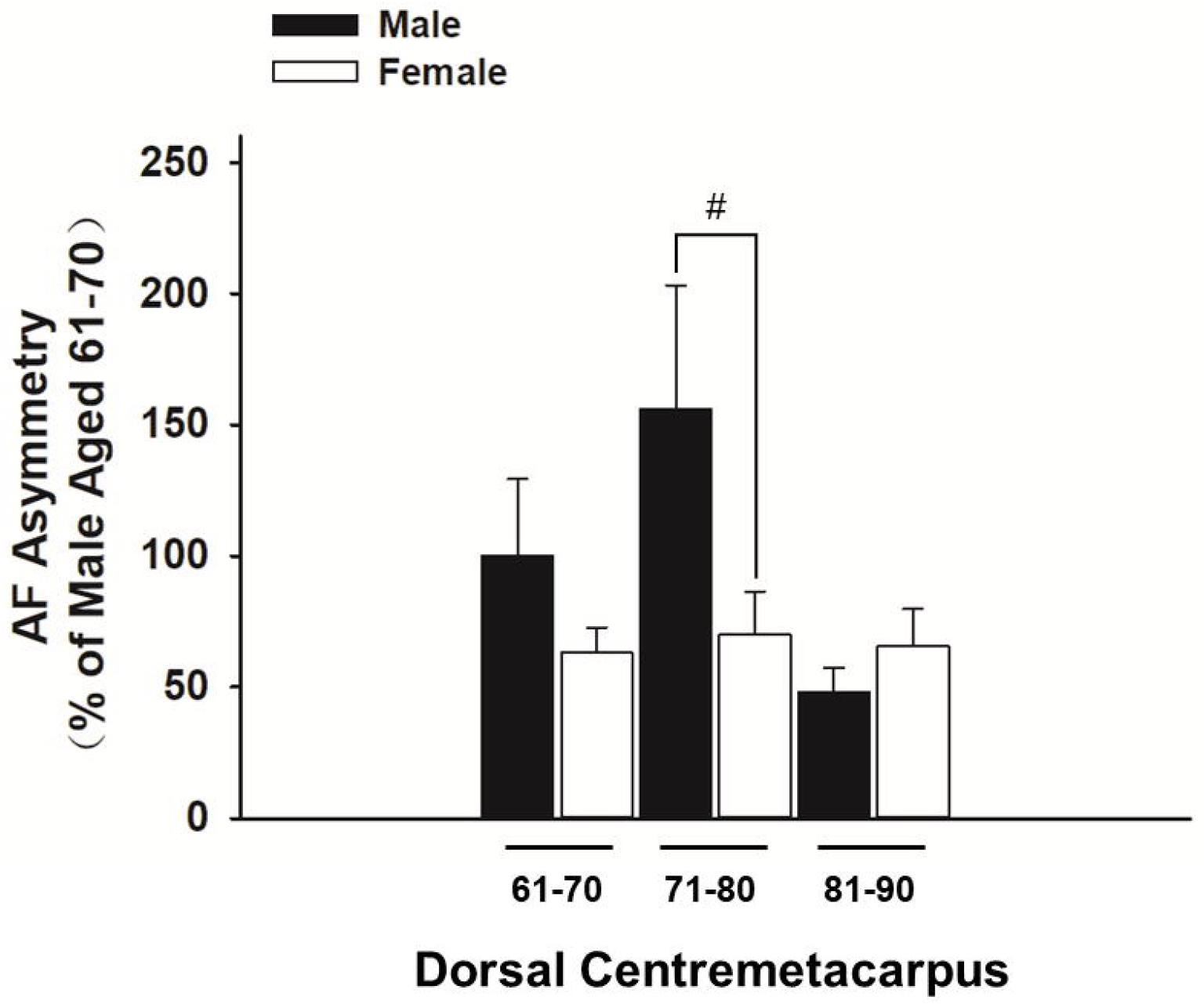
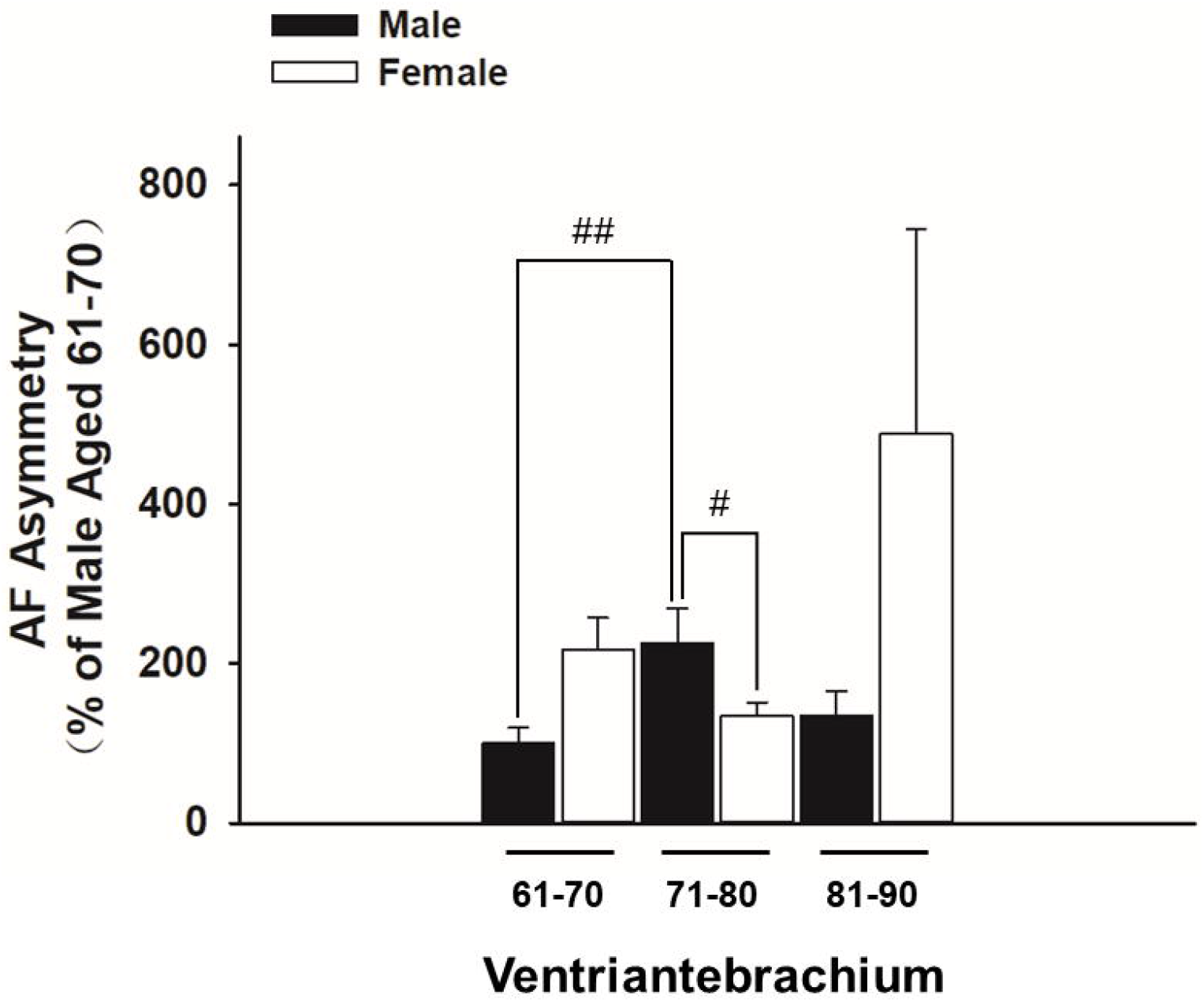
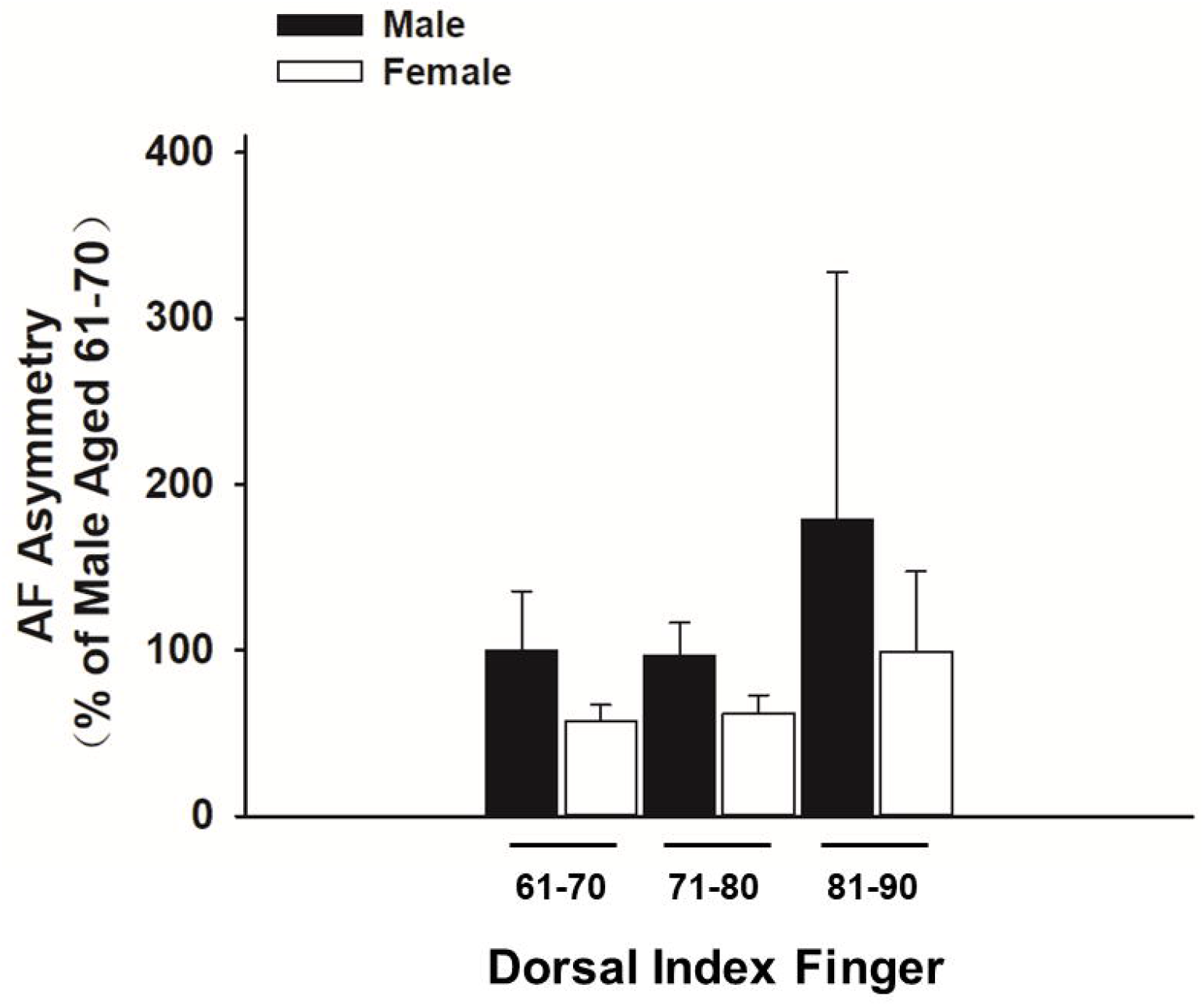
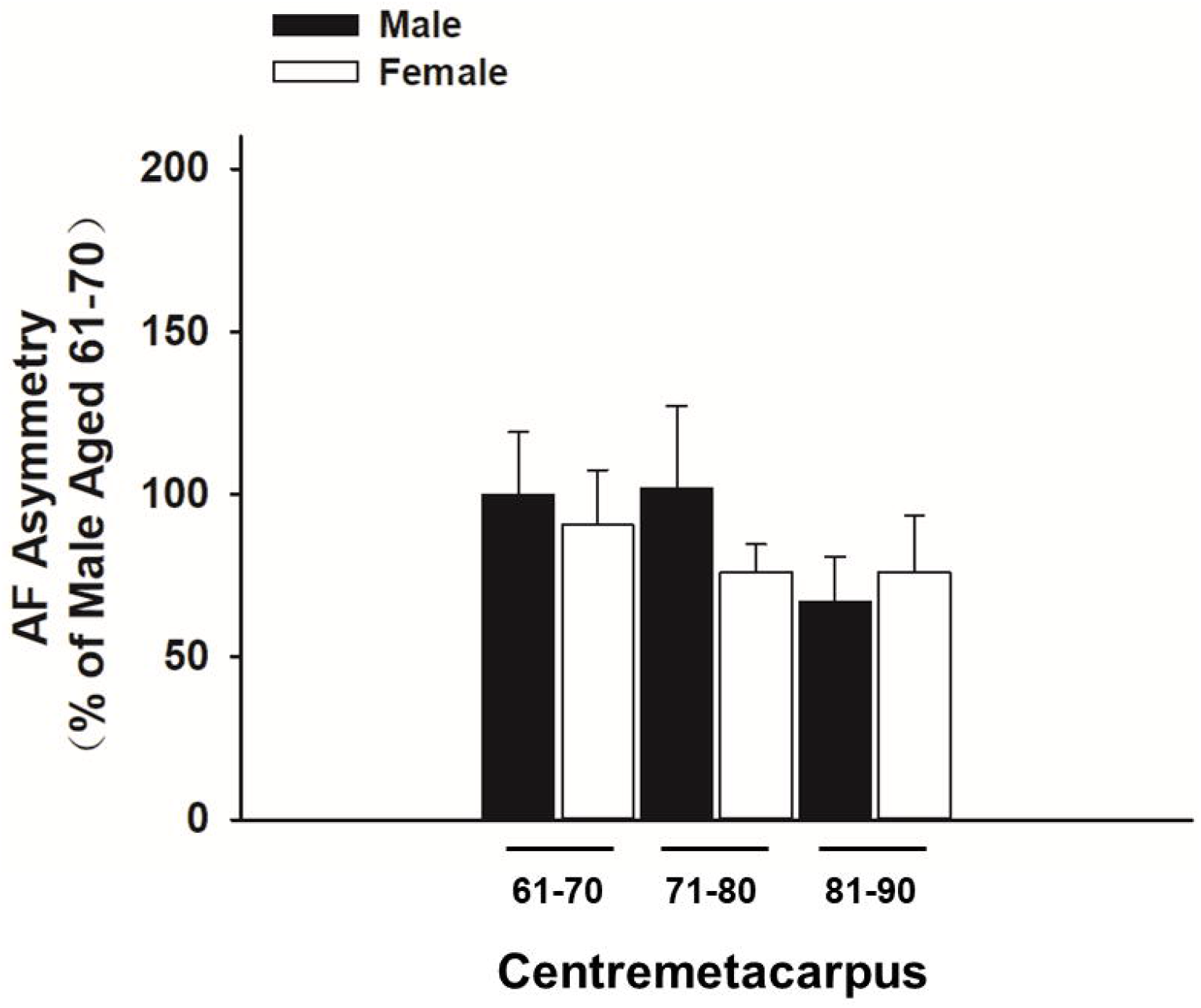
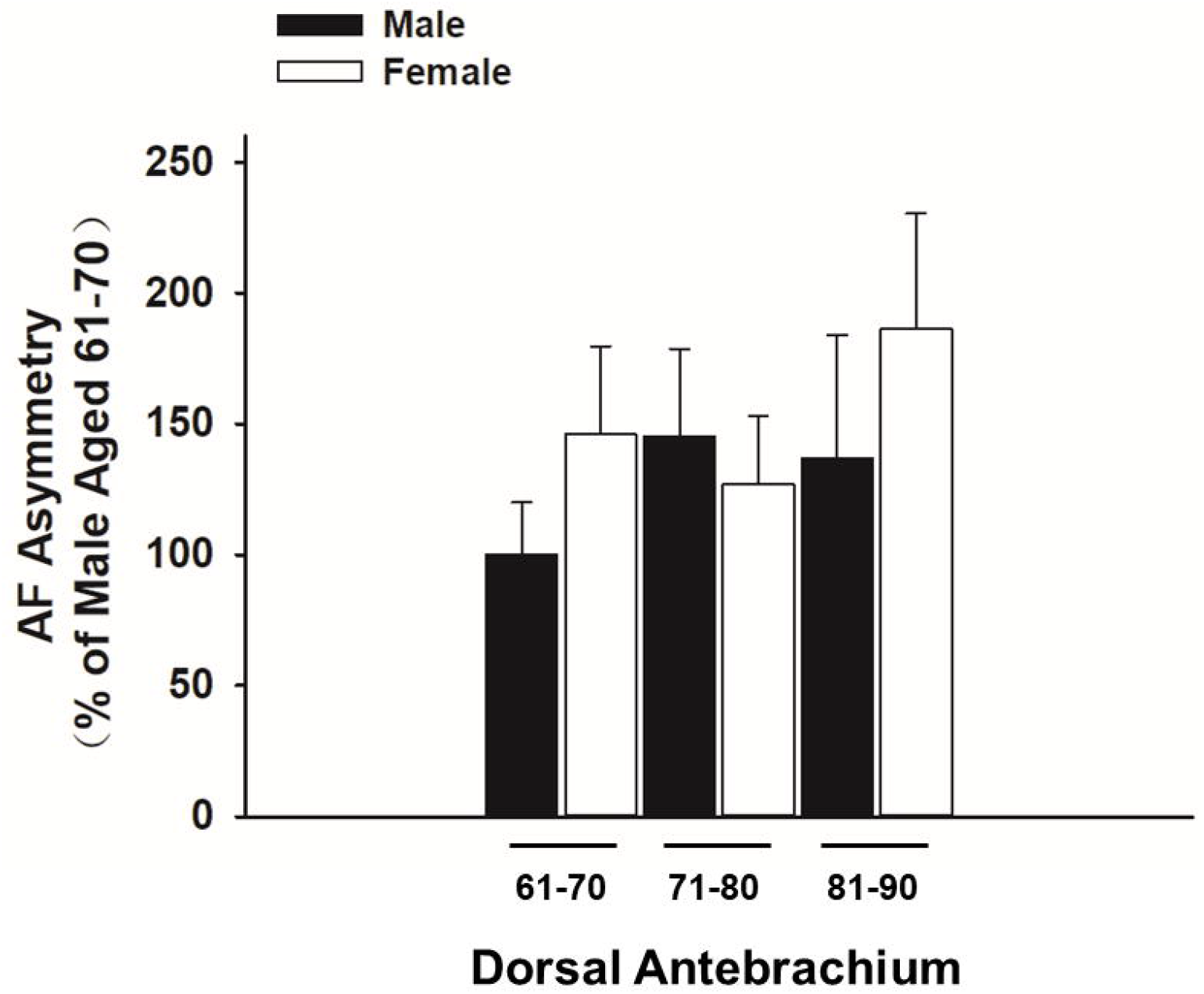
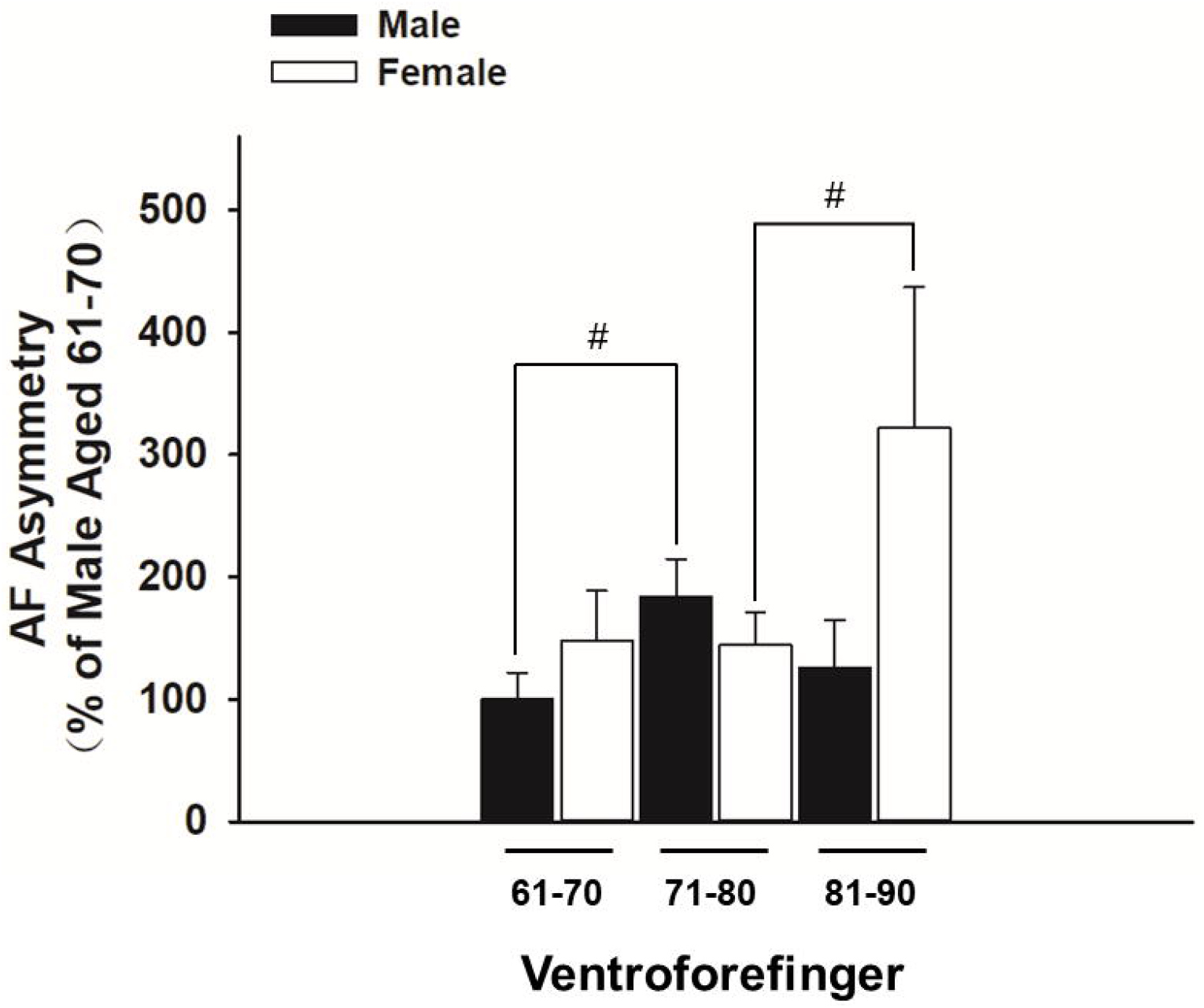
The green autofluorescence asymmetry of the men at the age between 71 - 80 years of old is significantly higher than that of the women of the same age group at both Dorsal Centremetacarpus and Centremetacarpus. (A) At Dorsal Centremetacarpus, the green AF asymmetry of the men at the age between 71 - 80 years of old is significantly higher than that of the women of the same age group. (B) At Centremetacarpus, we also found a similar difference between the green AF asymmetry of the men at the age between 71 - 80 years of old and that of the women of the same age group. At other regions examined, including Dorsal Index Finger (4C), Dorsal Antebrachium (4D), Ventroforefinger (4E) and Antebrachium (4F), there was no significant differences between the green AF asymmetry of the older men and that of the women of the same age groups. #, *P* < 0.05 (*t*-test); ##, *P* < 0.01 (*t*-test). The number of the men in the age group of 61 - 70, 71 - 80, and 81 - 90 years of old is 25, 20 and 11, respectively. The number of the women in the age group of 61 - 70, 71 - 80, and 81 - 90 years of old is 64, 48 and 15, respectively.

## Discussion

The major findings of our current study include: First, in both left and right Index Fingernails, the green AF intensity of the men of both the age group of 61 - 70 years of old and the age group of 71 - 80 years of old is significantly higher than that of the women of the same age groups; second, at both left Dorsal Centremetacarpus and left Centremetacarpus, the green AF intensity of the men at the age between 71 - 80 years of old is significantly higher than that of the women of the same age group; and third, in Index Fingernails, Dorsal Centremetacarpus and Centremetacarpus, the green AF asymmetry of the older men of certain age groups is significantly higher than that of the women of the same age groups. Collectively, our study has provided the first evidence indicating the gender difference between the green AF intensity and asymmetry of elderly men and elderly women in fingernails and certain regions of skin.

Our previous studies have suggested that each of the five major age-related diseases, including AIS (3), MI (17), stable coronary artery disease (17), Parkinson’s disease (4) and lung cancer (13), has its unique ‘Pattern of AF’. Numerous studies have reported that women have longer life expectancy than men (9), e.g., as reported by a study on the life expectancy of 54 countries from 1999 to 2004, on average women have 5 years longer life expectancy than men (15). The sex gap in life expectancy is also common in non-human animals, especially in mammals (9). Female also have significantly lower risk to develop multiple cardiovascular diseases including MI, hypertension and heart failure (7). It is noteworthy that the women who develop first MI could be 10 years older than the men who develop first MI (7).

Inflammation is a common key pathological factor of a number of age-related diseases (14). It has been reported that older women do not have decreased serum concentrations of IL-10, an anti-inflammatory cytokine, as older men do (1), which has implicated that older men may have higher levels of inflammation than older women. Since our previous study has suggested that inflammation can induce increased skin’s green AF intensity (19), the higher green AF intensity of older men’s skin may be caused by their higher levels of inflammation.

While several factors, including sex hormones, sex chromosomes, and the mother-based transmission of mitochondria, may contribute the sex gap of longevity (9), oxidative stress may play a significant role in the sex gap of longevity (7). Multiple studies have indicated that male have higher levels of oxidative stress than female (2,6,7,16). It has been reported that compared with elderly women, elderly men have significantly higher levels of DNA damage (8), suggesting higher levels of oxidative damage in elderly men: Oxidative stress is an important cause of DNA damage (18), particularly in aging body, as evidenced by the finding that there are approximately 24,000 and 66,000 steady state oxidative DNA adducts per cell in young and old rats, respectively (5). However, using homocysteine as a marker of oxidative stress, a study did not observe a significant difference between the post-menopause women and elderly men (7). Therefore, future studies are warranted to further determine if oxidative stress is an important factor accounting for the higher levels of the green AF intensity in older men.

Our previous study has indicated the green AF intensity is highly proportional to the risk of developing cardiovascular diseases (12). Therefore, our current finding that elder men have higher green AF intensity than elder women is consistent with the findings indicating that female have significantly lower risk to develop multiple cardiovascular diseases (7). Our studies have indicated that the green AF intensity is not only highly correlated with the risk to develop multiple cardiovascular diseases, but also a hallmark of tissue injury induced by oxidative stress or inflammation (10,11,19). Therefore, our study has suggested that compared with older female population, the older male population has higher injury levels in their cardiovascular systems.

To our knowledge, our current study has provided the first evidence indicating gender difference in the green AF of human’s fingernails and skin. AF has shown increasingly important value in biomedical applications. Since gender is a critical factor in aging and age-dependent diseases, our study indicating the gender difference of the green AF of human body may provide valuable information for establishing AF-based diagnostic approaches for multiple age-related diseases.

## Acknowledgment

The authors would like to acknowledge the financial support by a Major Special Program Grant of Shanghai Municipality (Grant # 2017SHZDZX01) (to W.Y.), and a Major Research Grant from the Scientific Committee of Shanghai Municipality #16JC1400500 and #16JC1400502 (to W.Y.).

